# Prioritization of genes driving congenital phenotypes of patients with *de novo* genomic structural variants

**DOI:** 10.1101/707430

**Authors:** Sjors Middelkamp, Judith M. Vlaar, Jacques Giltay, Jerome Korzelius, Nicolle Besselink, Sander Boymans, Roel Janssen, Lisanne de la Fonteijne, Ellen van Binsbergen, Markus J. van Roosmalen, Ron Hochstenbach, Daniela Giachino, Michael E. Talkowski, Wigard P. Kloosterman, Edwin Cuppen

## Abstract

**Background:** Genomic structural variants (SVs) can affect many genes and regulatory elements. Therefore, the molecular mechanisms driving the phenotypes of patients with multiple congenital abnormalities and/or intellectual disability carrying *de novo* SVs are frequently unknown.

**Results:** We applied a combination of systematic experimental and bioinformatic methods to improve the molecular diagnosis of 39 patients with *de novo* SVs and an inconclusive diagnosis after regular genetic testing. In seven of these cases (18%) whole genome sequencing analysis detected disease-relevant complexities of the SVs missed in routine microarray-based analyses. We developed a computational tool to predict effects on genes directly affected by SVs and on genes indirectly affected due to changes in chromatin organization and impact on regulatory mechanisms. By combining these functional predictions with extensive phenotype information, candidate driver genes were identified in 16/39 (41%) patients. In eight cases evidence was found for involvement of multiple candidate drivers contributing to different parts of the phenotypes. Subsequently, we applied this computational method to a collection of 382 patients with previously detected and classified *de novo SVs* and identified candidate driver genes in 210 cases (54%), including 32 cases whose SVs were previously not classified as pathogenic. Pathogenic positional effects were predicted in 25% of the cases with balanced SVs and in 8% of the cases with copy number variants.

**Conclusions:** These results show that driver gene prioritization based on integrative analysis of WGS data with phenotype association and chromatin organization datasets can improve the molecular diagnosis of patients with *de novo* SVs.

## Background

*De novo* germline structural variations (SVs) including deletions, duplications, inversions, insertions and translocations are important causes of (neuro-)developmental disorders such as intellectual disability and autism [1,2]. Clinical genetic centers routinely use microarrays and sometimes karyotyping to detect SVs at kilo- to mega base resolution [3]. The pathogenicity of an SV is generally determined by finding overlap with SVs in other patients with similar phenotypes [4,5]. SVs can affect large genomic regions which can contain many genes and non-coding regulatory elements [1]. This makes it challenging to determine which and how specific affected gene(s) and regulatory elements contributed to the phenotype of a patient. Therefore, the causative genes driving the phenotype are frequently unknown for patients with *de novo* SVs which can hamper conclusive genetic diagnosis.

SVs can have a direct effect on the expression and functioning of genes by altering their copy number or by truncating their coding sequences [1]. In addition, SVs can also indirectly influence the expression of adjacent genes by disrupting the interactions between genes and their regulatory elements [6]. New developments in chromatin conformation capture (3C) based technologies such as Hi-C have provided the means to study these indirect, positional effects [7]. Most of the genomic interactions (loops) between genes and enhancers occur within megabase-sized topologically associated domains (TADs). These domains are separated from each other by boundary elements characterized by CTCF-binding, which limit interactions between genes and enhancers that are not located within the same TAD [8,9]. For several loci, such as the *EPHA4* [10], *SOX9* [11], *IHH* [12], *Pitx* [13] loci, it has been demonstrated that disruption of TAD boundaries by SVs can cause rewiring of genomic interactions between genes and enhancers, which can lead to altered gene expression during embryonic development and ultimately in disease phenotypes [14]. Although the organization of TADs appears to be stable across cell types, sub-TAD genomic interactions between genes and regulatory elements have been shown to be relatively dynamic and cell type-specific [15]. Disruptions of genomic interactions are therefore optimally studied in disease-relevant cell types, which may be obtained from mouse models or from patient-derived induced pluripotent stem cells. However, it is not feasible to study each individual locus or patient with such elaborate approaches and disease-relevant tissues derived from patients are usually not available. Therefore, it is not yet precisely known how frequently positional effects contribute to the phenotypes of patients with developmental disorders.

It has been shown that the use of computational methods based on combining phenotypic information from the Human Phenotype Ontology (HPO) database (“phenomatching”) with previously published chromatin interactions datasets can improve the interpretation of the molecular consequences of *de novo* SVs [16–18]. These approaches have largely been based on data derived from a small set of cell types and techniques. Here, we further expand these *in silico* approaches by integrating detailed phenotype information with genome-wide chromatin conformation datasets of many different cell types. By combining this method with whole genome and transcriptome sequencing we predicted which genes are affected by the SVs and which of these genes have likely been involved in the development of the disease phenotype (e.g. candidate driver genes). Accurate characterization of the effects of SVs on genes can be beneficial for the prediction of potential clinical relevance of the SVs. Detailed interpretation of the molecular effects of the SVs helped to identify candidate driver genes in 16 out of 39 included patients who had an inconclusive diagnosis after regular genetic testing. By applying the computational method on larger cohorts of patients with *de novo* SVs we estimated the contribution of positional effects for both balanced and unbalanced SVs.

## Results

### WGS reveals hidden complexity of *de novo* SVs

We aimed to improve the genetic diagnosis of 39 individuals with multiple congenital abnormalities and/or intellectual disability (MCA/ID) who had an inconclusive diagnosis after regular genetic testing or who have complex genomic rearrangements. The phenotypes of the individuals were systematically described by Human Phenotype Ontology (HPO) terms [19–21]. The included individuals displayed a wide range of phenotypic features and most individuals (82%) presented neurological abnormalities including intellectual disability (Fig. 1a, Additional File 2: Table S1). The parents of each of the patients were healthy, suggesting a *de novo* or recessive origin of the disease phenotypes. All individuals carried *de novo* SVs which were previously detected by ArrayCGH, SNP arrays, karyotyping or long-insert mate-pair sequencing (Additional File 1: Fig. S1a). First, we performed whole genome sequencing (WGS) for all individuals in the cohort to screen for potential pathogenic genetic variants that were not detected by the previously performed genetic tests. No known pathogenic single nucleotide variants (SNVs) were detected in the individuals analysed by patient-parents trio-based WGS (individuals P1 to P20), except for one pathogenic SNV that is associated with a part (haemophilia) of the phenotype of individual P1. A total of 46 unbalanced and 219 balanced *de novo* SVs were identified in the genomes of the individuals (Fig.1b, Additional File 1: Fig. S1b, Additional File 2: Table S2). The detected SVs ranged from simple SVs to very complex genomic rearrangements that ranged from 4 to 40 breakpoint junctions per individual. Importantly, WGS confirmed all previously detected *de novo* SVs and revealed additional complexity of the SVs in 7 (39%) of the 18 cases who were not studied by WGS-based techniques before (Fig. 1c-d, Additional File 2: Table S2). In half of the cases with previously identified *de novo* copy number gains (4/8), the gains were not arranged in a tandem orientation, but instead they were inserted in another genomic region, which can have far-reaching consequences for accurate interpretation of the pathogenetic mechanisms in these individuals (Fig. 1d) [22–24]. This suggests that the complexity of copy number gains in particular is frequently underestimated by microarray analysis. For example, in one case (P11) a previously detected 170 kb copy number gain from chromosome 9 was actually inserted into chromosome X, 82 kb upstream of the *SOX3* gene (Fig. 1d, Additional File 1: Fig. S2). This inserted fragment contains a super-enhancer region that is active in craniofacial development[25] (Additional File 1: Fig. S2). The insertion of the super-enhancer may have disturbed the regulation of *SOX3* expression during palate development, which may have possibly caused the orofacial clefting in this individual [26–30]. The detection of these additional complexities in these seven patients exemplifies the added value that WGS analyses can have for cases that remain unresolved after standard array diagnostics [24].

**Fig. 1.**
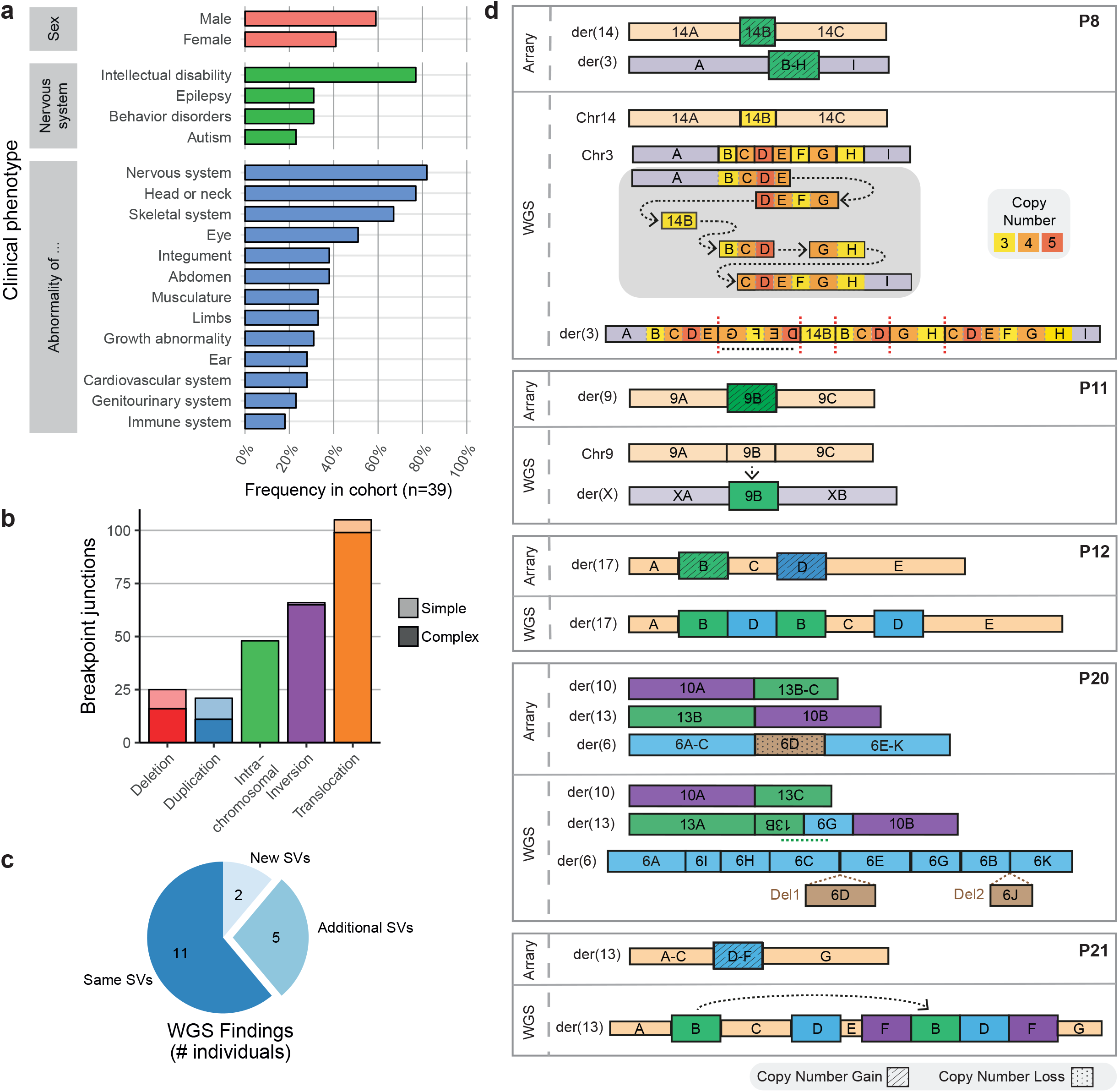
Characterization of *de novo* SVs in a cohort of individuals with neurodevelopmental disorders. **a** Frequencies of clinical phenotypic categories described for the 39 included individuals based on categories defined by HPO. Nervous system abnormalities are divided into four subcategories. **b** Number of *de novo* break junctions per SV type identified by WGS of 39 included patients. Most detected *de novo* SVs are part of complex genomic rearrangements involving more than three breakpoint junctions. **c** Number of cases in which WGS analysis identified new, additional or similar SVs compared to microarray-based copy number profiling. **d** Schematic representation of additional genomic rearrangements that were observed by WGS in five individuals. For each patient, the top panel shows the *de novo* SVs identified by arrays or karyotyping and bottom panel shows the structures of the SVs detected by WGS. The WGS data of individual P8 revealed complex chromoanasynthesis rearrangements involving multiple duplications and an insertion of a fragment from chr14 into chr3. Individual P11 has an insertion of a fragment of chr9 into chrX that was detected as a copy number gain by array-based analysis (Fig. S2). The detected copy number gains in individuals P12 and P21 show an interspersed orientation instead of a tandem orientation. The translocation in patient P20 appeared to be more complex than previously anticipated based on karyotyping results, showing 11 breakpoint junctions on three chromosomes.

### *In silico* phenomatching approach links directly affected genes to phenotypes

Subsequently, we determined if the phenotypes of the patients could be explained by direct effects of the *de novo* SVs, most of which were previously classified as a variant of unknown significance (VUS), on genes. In total, 332 genes are directly affected (deleted, duplicated or truncated) by the SVs in the cohort (Additional File 1: Fig. S1d). The Phenomatch tool was used to match the HPO terms associated with these genes with the HPO terms used to describe the phenotypes of the individuals [16,17]. Genes were considered as candidate driver genes based on the height of their Phenomatch score, the number of phenomatches between the HPO terms of the gene and the patient, recessive or dominant mode of inheritance, Loss of Function constraint score (pLI) [31], Residual Variation Intolerance Score (RVIS) [32] and the presence in OMIM and/or DD2GP [33] databases (Table 1). Directly affected genes strongly or moderately associated with the phenotype are classified as tier 1 (T1) and tier 2 (T2) candidate driver genes, respectively (Fig. 2a, Table 1). Genes with limited evidence for contribution to the phenotype are reported as tier 3 (T3) genes. In the cohort of 39 patients, this approach prioritized 2 and 13 of the 332 directly affected genes as T1 and T2 candidate drivers, respectively (Fig. 2b). In three cases, the HPO terms of the identified T1/T2 candidate driver genes could be matched to more than 75% of the HPO terms assigned to the patients, indicating that the effects of the SVs on these genes can explain most of the phenotypes of these patients. (Additional file 2: Table S3). In six other cases, directly affected T1/T2 candidate drivers were identified that were only associated with a part of the patient’s phenotypes (Additional file 2: Table S3).

**Fig. 2.**
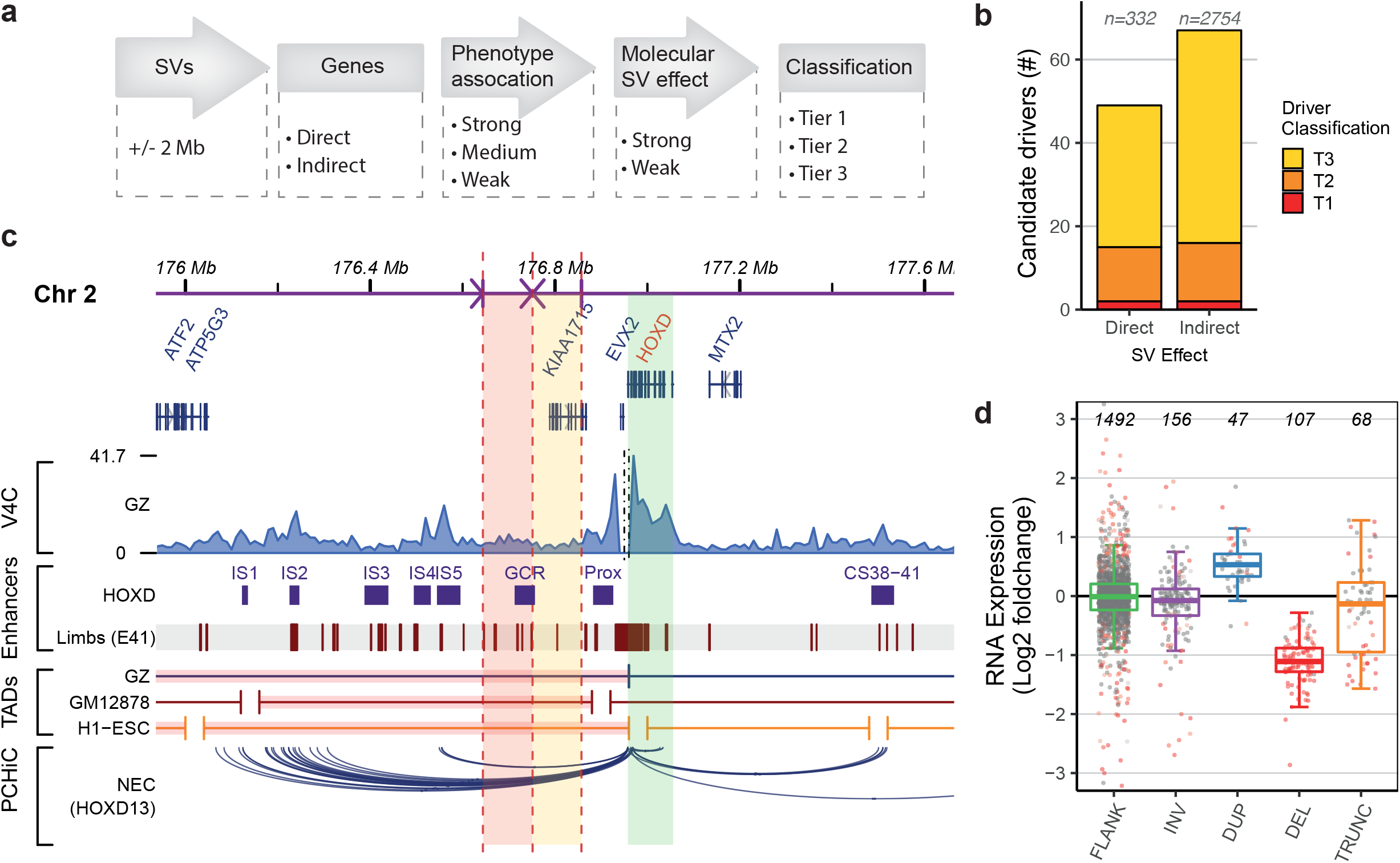
Prediction of candidate driver genes directly and indirectly affected by the SVs. **a** Schematic overview of the computational workflow developed to detect candidate driver genes. Classification of genes at (direct) or surrounding (indirect) the *de novo* SVs is based on association of the gene with the phenotype and the predicted direct or indirect effect on the gene (Table 1). **b** Total number of identified tier 1, 2 and 3 candidate driver genes predicted to be directly or indirectly affected by an SV. **c** Genome browser overview showing the predicted disruption of regulatory landscape of the *HOXD* locus in individual P22. A 107 kb fragment (red shading) upstream of the *HOXD* locus (green shading) is translocated to a different chromosome and a 106 kb fragment (yellow shading) is inverted. The SVs affect the TAD centromeric of the *HOXD* locus which is involved in the regulation of gene expression in developing digits. The translocated and inverted fragments contain multiple mouse [55] and human (day E41) [79] embryonic limb enhancers, including the global control region (GCR). Disruptions of these developmental enhancers likely contributed to the limb phenotype of the patient. The virtual V4C track shows the HiC interactions per 10kb bin in germinal zone (GZ) cells using the *HOXD13* gene as viewpoint [38]. The bottom track displays the PCHiC interactions of the *HOXD13* gene in neuroectodermal cells [42]. UCSC Liftover was used to convert mm10 coordinates to hg19. **d** RNA expression levels of genes at or adjacent to *de novo* SVs. Log2 fold RNA expression changes compared to controls (see methods) determined by RNA sequencing for expressed genes (RPKM >0.5) that are located within 2 Mb of SV breakpoint junctions (FLANK) or that are inverted (INV), duplicated (DUP), deleted (DEL) or truncated (TRUNC). Differentially expressed genes (p < 0.05, calculated by DESeq2) are displayed in red.

**Table 1:**
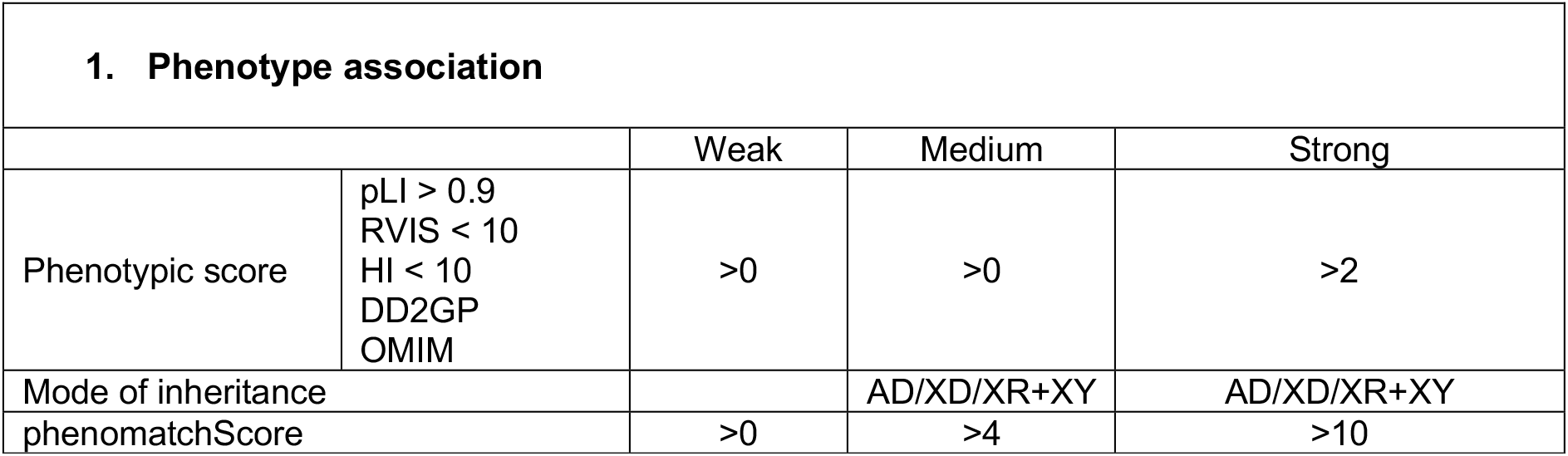
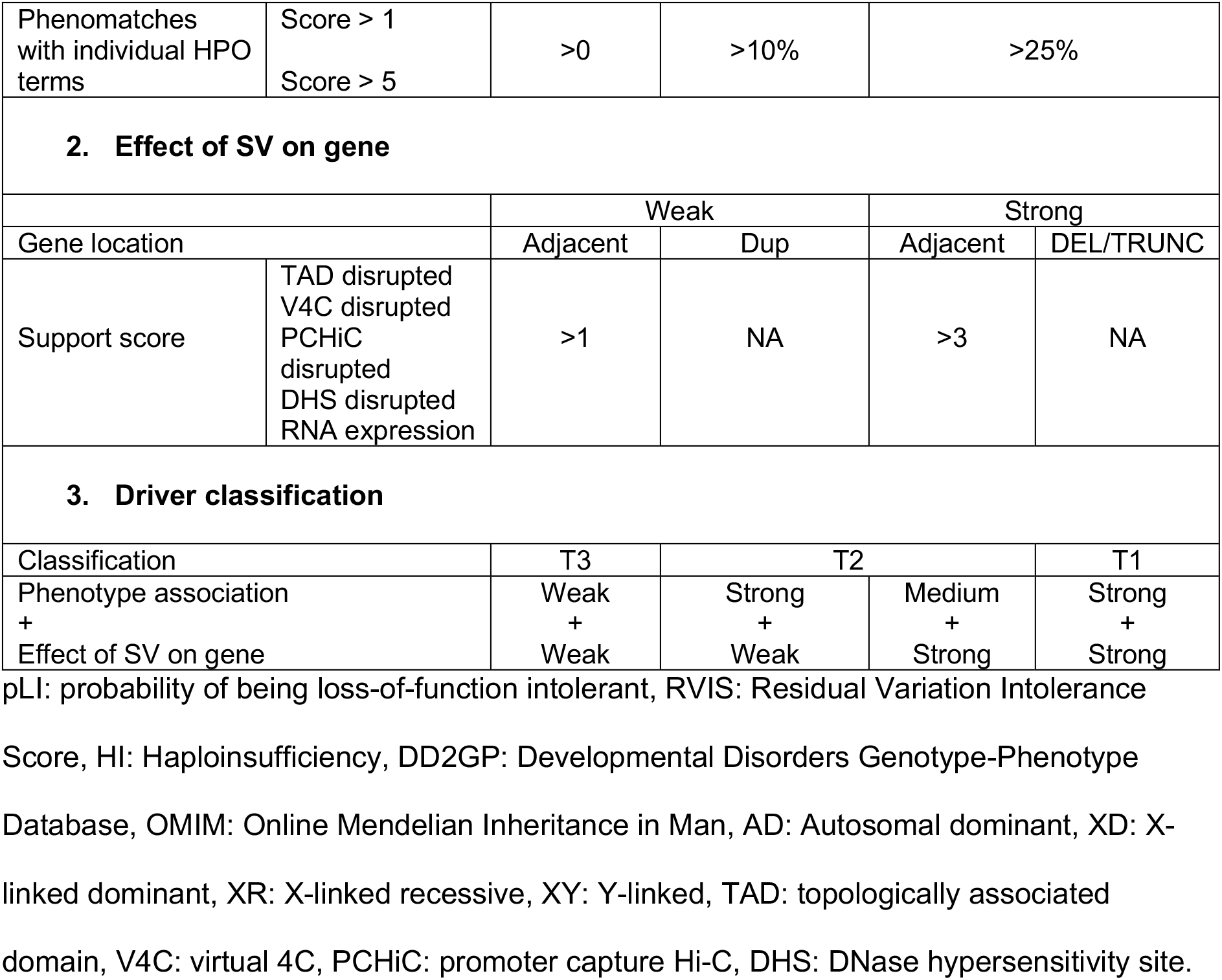
Cut-offs used to classify affected genes as T1, T2 or T3 candidate driver genes.

| 1. Phenotype association |  |  |  |  |
| --- | --- | --- | --- | --- |
|  |  | Weak | Medium | Strong |
| Phenotypic score | pLI > 0.9<br>RVIS < 10<br>HI < 10<br>DD2GP<br>OMIM | >0 | >0 | >2 |
| Mode of inheritance |  |  | AD/XD/XR+XY | AD/XD/XR+XY |
| phenomatchScore |  | >0 | >4 | >10 |

|  |  |  |  |  |  |
| --- | --- | --- | --- | --- | --- |
| Phenomatches with individual HPO terms | Score > 1<br>Score > 5 | >0 | >10% | >25% |  |
| 2. Effect of SV on gene |  |  |  |  |  |
|  |  | Weak |  | Strong |  |
| Gene location |  | Adjacent | Dup | Adjacent | DEL/TRUNC |
| Support score | TAD disrupted<br>V4C disrupted<br>PCHiC disrupted<br>DHS disrupted<br>RNA expression | >1 | NA | >3 | NA |
| 3. Driver classification |  |  |  |  |  |
| Classification |  | T3 | T2 |  | T1 |
| Phenotype association +<br>Effect of SV on gene |  | Weak +<br>Weak | Strong +<br>Weak | Medium +<br>Strong | Strong +<br>Strong |

pLI: probability of being loss-of-function intolerant, RVIS: Residual Variation Intolerance
Score, HI: Haploinsufficiency, DD2GP: Developmental Disorders Genotype-Phenotype
Database, OMIM: Online Mendelian Inheritance in Man, AD: Autosomal dominant, XD: X-
linked dominant, XR: X-linked recessive, XY: Y-linked, TAD: topologically associated
domain, V4C: virtual 4C, PCHiC: promoter capture Hi-C, DHS: DNase hypersensitivity site.

Subsequently, we performed RNA sequencing on primary blood cells or lymphoblastoid cell lines derived from the individuals to determine the impact of *de novo* SVs on RNA expression of candidate driver genes. RNA sequencing confirmed that most expressed genes directly affected by *de novo* deletions show a reduced RNA expression (97 of 107 genes with a median reduction of 0.46-fold compared to non-affected individuals) (Fig. 2d). Although duplicated genes show a median 1.44-fold increase in expression, only 14 of 43 (~30%) of them are significantly overexpressed compared to expression levels in non-affected individuals. In total, 87 genes are truncated by SVs and four of these are classified as T1/T2 candidate drivers. The genomic rearrangements lead to 12 possible fusions of truncated genes and RNA-seq showed an increased expression for two gene fragments due to formation of a fusion gene (Additional file 1: Fig. S3, Additional file 2: Table S4). However, none of the genes involved in the formation of fusion genes were associated with the phenotypes of the patients, although we cannot exclude an unknown pathogenic effect of the newly identified fusion genes. We could detect expression for three deleted and two duplicated T1/T2 candidate drivers and these were differentially expressed when compared to controls. The RNA sequencing data suggests that most genes affected by *de novo* deletions show reduced RNA expression levels and limited dosage compensation. However, the observed increased gene dosage by *de novo* duplications does not always lead to increased RNA expression, at least in blood cells of patients.

### Prediction of positional effects of *de novo* SVs on neighbouring genes

In 28 of the included cases (72%) our prioritization method did not predict T1/T2 candidate driver genes that are directly affected by the *de novo* SVs. Therefore, we investigated positional effects on the genes surrounding the *de novo* SVs to explain the phenotypes in those cases that were not fully explained by directly affected candidate driver genes. We extended our candidate driver gene prioritization analysis by including all the protein-coding genes located within 2 Mb of the breakpoint junctions, as most chromatin interactions are formed between loci that are less than 2Mb apart from each other [34]. Of the 2,754 genes adjacent to the SVs, 117 are moderately to strongly associated with the specific phenotypes of the individuals based on phenotype association analysis. However, this association with the phenotype does not necessarily mean that these genes located within 2Mb of the breakpoint junctions are really affected by the SVs and thus contributing to the phenotype. To determine if the regulation of these genes was affected, we first evaluated RNA expression levels of those genes. Three-quarters (81/117) of the genes linked to the phenotypes were expressed, but only 9 of these showed reduced or increased expression (Fig. 2d). However, RNA expression in blood may not always be a relevant proxy for most neurodevelopmental phenotypes [35,36]. Therefore, we developed an extensive *in silico* strategy to predict potential disruption of the regulatory landscape of the genes surrounding the SVs (Additional file 1: Fig. S4). Because the interactions between genes and their regulatory elements are cell-type specific a large collection of tissue-specific Hi-C, TAD, promoter capture Hi-C (PCHiC), DNase hypersensitivity site (DHS), RNA and ChIP-seq datasets was included (Additional file 2: Table S5). Several embryonic and neural cell type (such as fetal brain and neural progenitor cells) datasets were included that may be especially relevant to study the neurodevelopmental phenotypes in our cohort.

For each gene, we selected the TAD it is located in [37–39], the PCHiC interactions with its transcription start site [40–43] and its DHS connections [44] for each assessed cell type and overlapped these features with the breakpoint junctions of the SVs to determine the proportion of disrupted genomic interactions (Methods, Additional file 1: Fig. S4). We also counted the number of enhancers (which are active in cell types in which the genes show the highest RNA expression [45]) that are located on disrupted portions of the TADs. Additionally, we performed virtual 4C (v4C) for each gene by selecting the rows of the normalized Hi-C matrices containing the transcription start site coordinates of the genes as viewpoints, because the coordinates of TAD boundaries can be dependent on the calling method and the resolution of the Hi-C [46–48] and because a significant portion of genomic interactions crosses TAD boundaries [9]. Integrated scores for TAD disruption, v4C disruption, potential enhancer loss, disruption of PCHiC interactions and DHS connections were used to calculate a positional effect support score for each gene (Additional file 1: Fig. S4). Finally, indirectly affected genes were classified as tier 1, 2 or 3 candidate drivers based on a combination of their association with the phenotype and their support score (Fig. 2a, Table 1).

Of the 117 genes that were associated with the phenotypes and located within 2 Mb of the SVs, 16 genes were predicted to be affected by the SVs based on the *in silico* analysis and therefore classified as T1/T2 candidate driver gene (Fig. 2b). The validity of the approach was supported by the detection of pathogenic positional effects identified in previous studies. For example, the regulatory landscape of *SOX9* was predicted to be disturbed by a translocation 721 Kb upstream of the gene in individual P5, whose phenotype is mainly characterized by acampomelic campomelic dysplasia with Pierre-Robin Syndrome (PRS) including a cleft palate (Additional file 1: Fig. S6). SVs in this region have been predicted to disrupt interactions of *SOX9* with several of its enhancers further upstream, leading to phenotypes similar to the phenotype of individual P5 [49,50]. In individual P39, who has been previously included in other studies, our method predicted a disruption of *FOXG1* expression regulation due to a translocation (Additional file 1: Fig. S4), further supporting the hypothesis that deregulation of *FOXG1* caused the phenotype of this individual [51,52].

Another example of a predicted positional effect is the disruption of the regulatory landscape of the *HOXD* locus in individual P22. This individual has complex genomic rearrangements consisting of 40 breakpoint junctions on four different chromosomes likely caused by chromothripsis [53]. One of the inversions and one of the translocations are located in the TAD upstream (centromeric) of the *HOXD* gene cluster (Fig. 2c). This TAD contains multiple enhancers that regulate the precise expression patterns of the *HOXD* genes during the development of the digits [54–56]. Deletions of the gene cluster itself, but also deletions upstream of the cluster are associated with hand malformations [57–59]. The translocation in individual P22 disrupts one of the main enhancer regions (the global control region (GCR)), which may have led to altered regulation of the expression of *HOXD* genes, ultimately causing brachydactyly and clinodactyly in this patient.

Our approach predicted positional effects on T1/T2 candidate driver genes in 10 included cases (26%). Most of these predicted positional effects were caused by breakpoint junctions of balanced SVs, suggesting that these effects may be especially important for balanced SVs.

### Prediction of driver genes improves molecular diagnosis

By combining both directly and indirectly affected candidate drivers per patient we found possible explanations for the phenotypes of 16/39 (41%) complex and/or previously unsolved cases (Fig. 3a, Additional file 2: Table S3). Interestingly, in eight cases we found evidence for multiple candidate drivers that are individually only associated with part of the phenotype, but together may largely explain the phenotype (Fig. 3b). For example, we identified four candidate drivers in individual P25, who has a complex phenotype characterized by developmental delay, autism, seizures, renal agenesis, cryptorchidism and an abnormal facial shape (Fig. 3c). This individual has complex genomic rearrangements consisting of six breakpoint junctions and two deletions of ~10 Mb and ~0.6 Mb on three different chromosomes (Fig. 3d). The 6q13q14.1 deletion of ~10 Mb affects 33 genes including the candidate drivers *PHIP* and *COL12A1,* which have been associated with developmental delay, anxiety and facial dysmorphisms in other patients [60,61]. In addition to the two deleted drivers, two genes associated with other parts of the phenotype were predicted to be affected by positional effects (Fig. 3e). One of these genes is *TFAP2A*, whose TAD (characterized by a large gene desert) and long-range interactions overlap with a translocation breakpoint junction. Rearrangements affecting the genomic interactions between *TFAP2A* and enhancers active in neural crest cells located in the *TFAP2A* TAD have recently been implicated in branchio-oculofacial syndrome [62]. The regulation of *BMP2*, a gene linked to agenesis of the ribs and cardiac features, is also predicted to be disturbed by a complex SV upstream of this gene [63,64]. Altogether, these candidate driver genes may have jointly contributed to the phenotype of this individual (Fig. 3d). This case illustrates the challenge of identifying the causal genes driving the phenotypes of patients with structural rearrangements and highlights the notion that multiple genes should be considered for understanding the underlying molecular processes and explaining the patient’s phenotype.

**Fig. 3.**
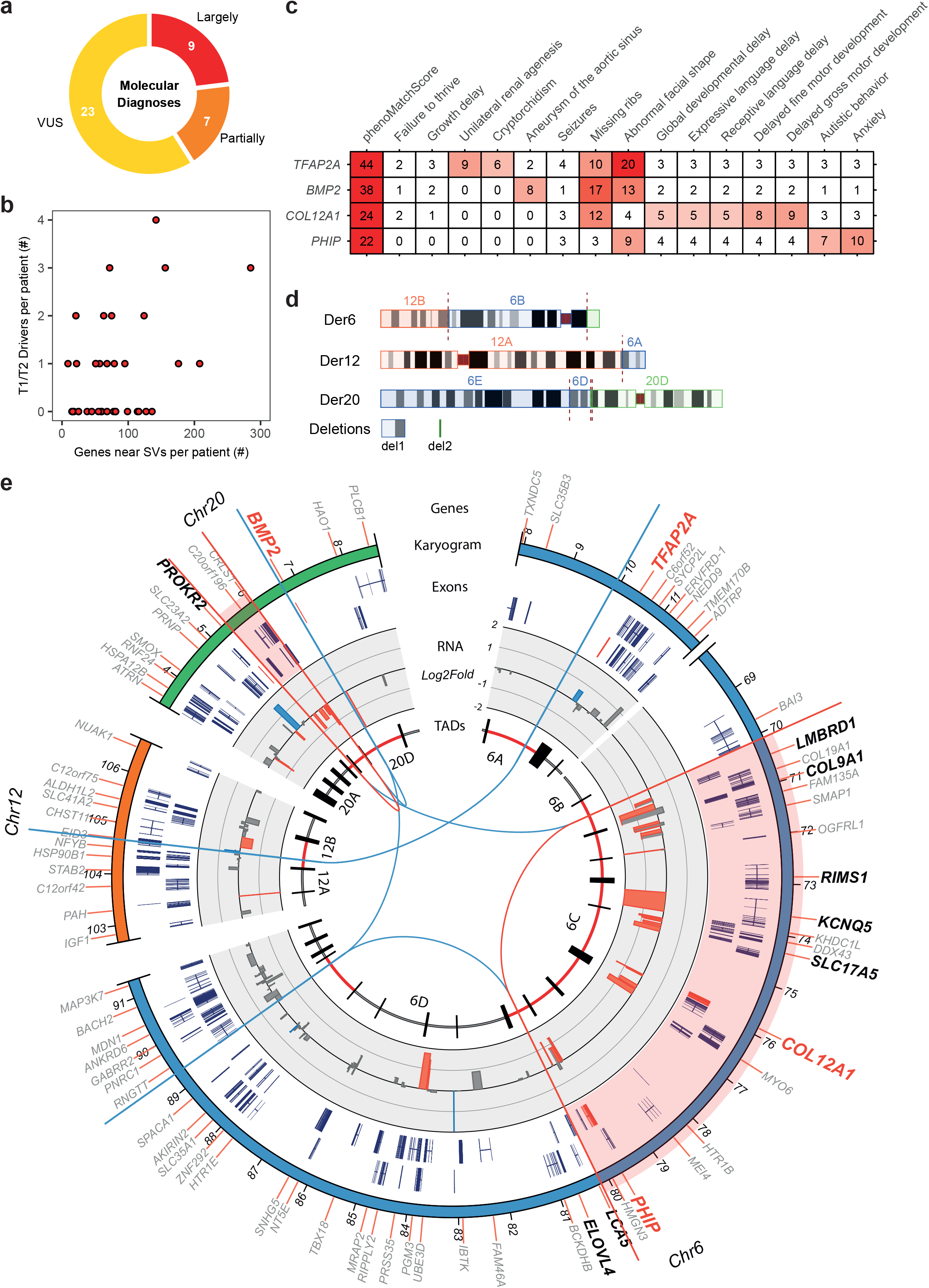
SVs can affect multiple candidate drivers which jointly contribute to a phenotype. **a** Number of patients whose phenotype can be partially or largely explained by the predicted T1/T2 candidate drivers. These molecular diagnoses are based on the fraction of HPO terms assigned to the patients that have a phenomatchScore of more than five with at least one T1/T2 driver gene. **b** Scatterplot showing the number of predicted T1/T2 candidate drivers compared to the total number of genes at or adjacent (< 2Mb) to the *de novo* SVs per patient. **c** Heatmap showing the association of the four predicted T1/T2 candidate drivers with the phenotypic features (described by HPO terms) of individual P25. The numbers correspond to score determined by Phenomatch. The four genes are associated with different parts of the complex phenotype of the patient. **d** Ideogram of the derivative (der) chromosomes 6, 12 and 20 in individual P25 reconstructed from the WGS data. WGS detected complex rearrangements with six breakpoint junctions and two deletions on chr6 and chr20 of respectively ~10 Mb and ~0.6 Mb. **e** Circos plot showing the genomic regions and candidate drivers affected by the complex rearrangements in individual P25. Gene symbols of T1/T2 and T3 candidate drivers are shown in respectively red and black. The break junctions are visualized by the lines in the inner region of the plot (red lines and highlights indicate the deletions). The middle ring shows the log2 fold change RNA expression changes in lymphoblastoid cells derived from the patient compared to controls measured by RNA sequencing. Genes differentially expressed (p<0.05) are indicated by red (log2 fold change < −0.5) and blue (log2 fold change > 0.5) bars. The inner ring shows the organization of the TADs and their boundaries (indicated by vertical black lines) in germinal zone (GZ) brain cells [38]. TADs overlapping with the *de novo* SVs are highlighted in red.

### *In silico* driver gene prediction in larger patient cohorts

Our candidate driver prioritization approach identified many candidate drivers in previously unresolved cases, but these complex cases may not be fully representative for the general patient population seen in clinical genetic diagnostics. Therefore, we applied our prediction method to two larger sets of patients with *de novo* SVs to further assess the validity and value of the approach. We focused on the genes located at or within 1 Mb of the SVs, because most of the candidate driver genes we identified in our own patient cohort were located within 1 Mb of an SV breakpoint junction (Additional file 1: Fig. S5b). First, we determined the effects of largely balanced structural variants in 228 previously described patients (Additional file 1: Fig. S7a) [51]. In 101 of the 228 (44%) cases the detected *de novo* SVs were previously classified as pathogenic or likely pathogenic and in all but four of these diagnosed cases one or more candidate driver genes have been proposed (Additional file 1: Fig. S7b). Our approach identified 46 T1 and 92 T2 candidate drivers out of 7406 genes located within 1 Mb of the SVs (Additional file 1: Fig. S7c,d, Additional file 2: Table S7). More than half (85/138) of the identified T1/T2 candidate drivers were not previously described as driver genes. In contrast, 23/114 (22%) previously described pathogenic or likely pathogenic drivers were classified as T3 candidates and 38/114 (33%) were not reported as driver by our approach (Fig. 4a), mostly because the Phenomatch scores were below the threshold (46%) or because the genes were not associated with HPO terms (41%) (Additional file 1: Fig. S7e). T1/T2 candidate drivers were identified in 99/225 (44%) of the individuals with mostly balanced SVs, including 31 individuals with SVs that were previously classified as VUS (Fig. 4b, Additional file 1: Fig. S8). Positional effect on genes moderately to strongly associated with the phenotypes were predicted in 63 (28%) of the cases with balanced SVs.

**Fig. 4.**
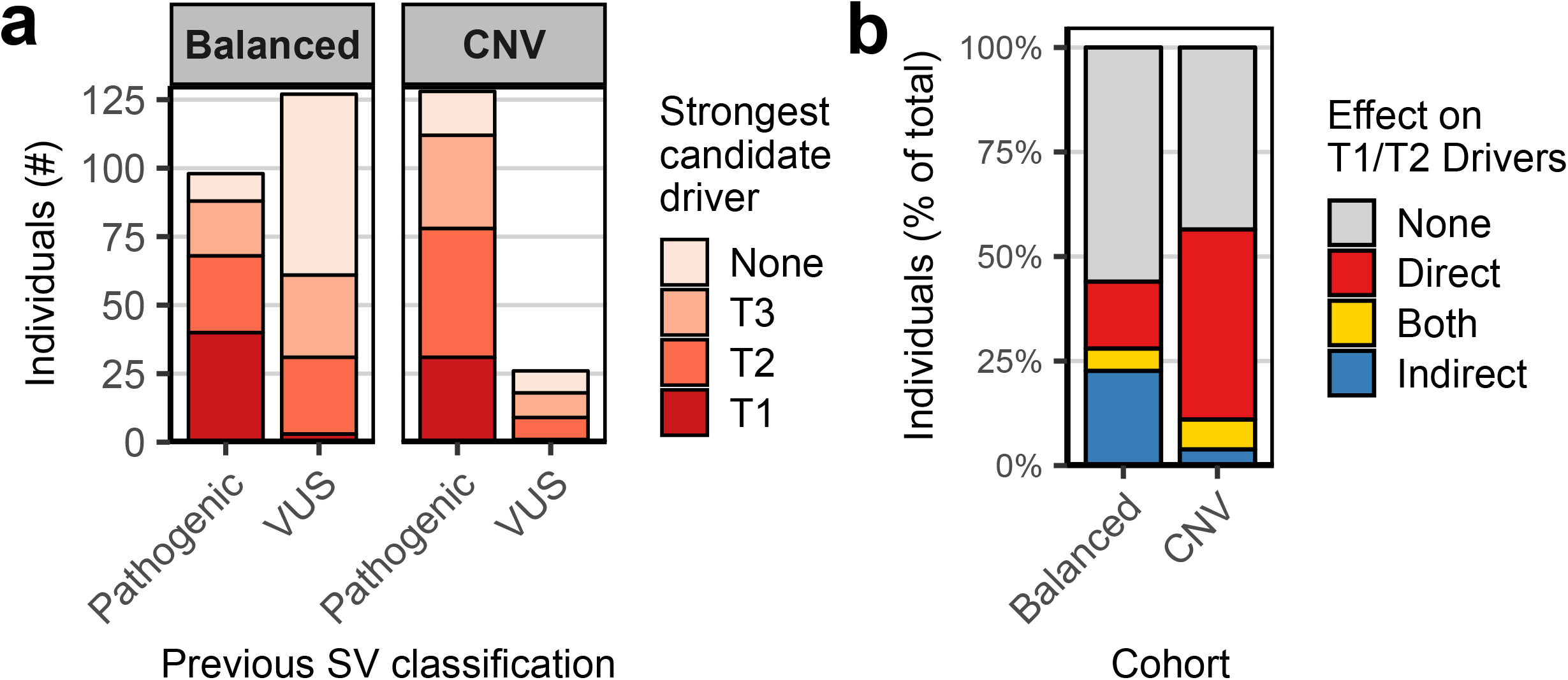
*In silico* prediction of candidate drivers in larger cohorts of patients with *de novo* SVs. **a** Comparison between previous SV classifications with the strongest candidate driver (located at or adjacent (<1 Mb) to these SVs) predicted by our approach. Two different patient cohorts, one containing mostly balanced SVs [51] and one containing copy number variants, were screened for candidate drivers. Our method identified T1/T2 candidate drivers for most SVs previously classified as pathogenic or likely pathogenic. Additionally, the method detected T1/T2 candidate drivers for some SVs previously classified as VUS, which may lead to a new molecular diagnosis. **b** Quantification of the predicted effects of the SVs on proposed T1/T2 candidate driver genes per cohort. Individuals with multiple directly and indirectly affected candidate drivers are grouped in the category described as “Both”. Indirect positional effects of SVs on genes contributing to phenotypes appear to be more common in patients with balanced SVs compared to patients with copy number variants.

Subsequently, we also assessed the value of our driver prioritization approach for individuals with unbalanced copy number variants. We collected genetic and phenotypic information of 154 individuals with *de novo* copy number variants (<10 Mb) identified by clinical array-based copy number profiling (Additional file 1: Fig. S7a, b, Additional file 2: Table S6). The CNVs in the majority (83%) of these individuals have been previously classified as pathogenic according to clinical genetic diagnostic criteria (Additional file 1: Fig. S7b). These criteria are mostly based on the overlap of the CNVs with CNVs of other individuals with similar phenotypes and the causative driver genes were typically not previously specified. Our method identified T1/T2 candidate driver genes in 87/154 (56%) individuals, including 9/26 individuals with CNVs previously classified as VUS (Fig. 4a, Additional file 2: Table S7). Interestingly, support for positional effects on candidate drivers was only found in 11% of the cases with CNVs, suggesting that pathogenic positional effects are more common in patients with balanced SVs than in patients with unbalanced SVs (Fig. 4b). No driver genes were identified for 39% of the previously considered pathogenic CNVs (based on recurrence in other patients). In some cases, potential drivers may remain unidentified because of incompleteness of the HPO database or insufficient description of the patient’s phenotypes. However, given the WGS results described for our patient cohort, it is also likely that some complexities of the CNVs may have been missed by the array-based detection method. The data also suggests that many disease-causing genes or mechanisms are still not known, and that some SVs are incorrectly classified as pathogenic.

## Discussion

About half of the patients with neurodevelopmental disorders do not receive a diagnosis after regular genetic testing based on whole exome sequencing and microarray-based copy number profiling [3]. Furthermore, the molecular mechanisms underlying the disease phenotype often remain unknown, even when a genetic variant is diagnosed as (potentially) pathogenic in an individual, as this is often only based on recurrence in patients with a similar phenotype. Here, we applied an integrative method based on WGS, computational phenomatching and prediction of positional effects to improve the diagnosis and molecular understanding of disease aetiology of individuals with *de novo* SVs.

Our WGS-approach identified additional complexities of the *de novo* SVs previously missed by array-based analysis in 7 of 18 cases, supporting previous findings that WGS can have an added value in identifying additional SVs that are not routinely detected by microarrays [24]. Our results indicate that duplications in particular are often more complex than interpreted by microarrays, which is in line with previous studies [22,65]. WGS can therefore be a valuable follow-up method to improve the diagnosis particularly of patients with copy number gains classified as VUS. Knowing the exact genomic location and orientation of SVs is important for the identification of possible positional effects.

To systematically dissect and understand the impact of *de novo* SVs, we developed a computational tool based on integration of HiC, RNA-seq and ChIP-seq datasets to predict positional effects of SVs on the regulation of gene expression. We combined these predictions with phenotype association information to identify candidate driver genes. In 9/39 of the complex cases we identified candidate drivers that are directly affected by the break-junctions of the SVs. Positional effects of SVs have been shown to cause congenital disorders, but their significance is still unclear and they are not yet routinely screened for in genetic diagnostics [14]. Our method predicted positional effects on genes associated with the phenotype in 25% and 8% of all studied cases with balanced and unbalanced *de novo* SVs, respectively. Previous studies estimated that disruptions of TAD boundaries may be the underlying cause of the phenotypes of ~7.3% patients with balanced rearrangements [51] and of ~11.8% of patients with large rare deletions [16]. Our method identified a higher contribution of positional effects in patients with balanced rearrangements mainly because our method included more extensive chromatin conformation datasets and also screened for effects that may explain smaller portions of the phenotypes. Our method, although it incorporates most of all published chromatin conformation datasets on untransformed human cells, focuses on disruptions of interactions, which is a simplification of the complex nature of positional effects. It gives an insight in the potential effects that lead to the phenotypes and prioritizes candidates that need to be followed up experimentally, ideally in a developmental context for proofing causality.

SVs can affect many genes and multiple ‘disturbed’ genes may together contribute to the phenotype. Indeed, in eight cases we found support for the involvement of multiple candidate drivers that were affected by one or more *de novo* SVs. Nevertheless, in many of the studied cases our method did not detect candidate drivers. This may be due to insufficient data or knowledge about the genes and regulatory elements in the affected locus and/or due to missing disease associations in the used databases. Additionally, *de novo* SVs are also frequently identified in healthy individuals [66–68] and some of the detected SVs of unknown significance may actually be benign and the disease caused by other genetic or non-genetic factors. The datasets underlying our computational workflow can be easily updated with more detailed data when emerging in the future, thereby enabling routine reanalysis of previously identified SVs. Moreover, our approach can be extended to study the consequences of SVs in different disease contexts such as cancer, where SVs also play a major causal role.

## Conclusions

Interpretation of SVs is important for clinical diagnosis of patients with developmental disorders, but it remains a challenge because SVs can have many different effects on multiple genes. We developed an approach to gain a detailed overview of the genes and regulatory elements affected by *de novo* SVs in patients with congenital disease. We show that WGS can be useful as a second-tier test to detect variants that are not detected by exome- and array-based approaches.

## Methods

### Patient selection and phenotyping

A total of 39 individuals with *de novo* germline SVs and an inconclusive diagnosis were included in this study. Individuals P1 to P21 and their biological parents were included at the University Medical Center Utrecht (the Netherlands) under study ID NL55260.041.15 15-736/M. Individual P22, previously described by Redin *et al.* as UTR22 [51], and her parents were included at the San Luigi University Hospital (Italy). For individuals P23 to P39, lymphoblastoid cell lines (LCL cell lines) were previously derived as part of the Developmental Genome Anatomy Project (DGAP) of the Brigham and Women’s Hospital and Massachusetts General Hospital, Boston, Massachusetts, USA [51]. Written informed consent was obtained for all included individuals and parents and the studies were approved by the respective institutional review boards.

### DNA and RNAextraction

Peripheral blood mononuclear cells (PBMCs) were isolated from whole blood samples of individuals P1 to P22 and their biological parents using a Ficoll-Paque Plus gradient (GE Healthcare Life Sciences) in SepMate tubes (STEMCELL Technologies) according to the manufacturer’s protocols. LCL cell lines derived from individuals P23 to P39 were expanded in RPMI 1640 medium supplemented with GlutaMAX (ThermoFisher Scientific), 10% fetal bovine serum, 1% penicillin, 1% streptomycin at 37°C. LCL cultures of each individual were split into three flasks and cultured separately for at least one week to obtain technical replicate samples for RNA isolation. Genomic DNA was isolated from the PBMCs or LCL cell lines using the QIASymphony DNA kit (Qiagen). Total RNA was isolated using the QIAsymphony RNA Kit (Qiagen) and RNA quality (RIN > 8) was determined using the Agilent RNA 6000 Nano Kit.

### Whole-genome sequencing

Purified DNA was sheared to fragments of 400-500 bp using a Covaris sonicator. WGS libraries were prepared using the TruSeq DNA Nano Library Prep Kit (Illumina). WGS libraries were sequenced on an Illumina Hiseq X instrument generating 2×150 bp paired-end reads to a mean coverage depth of at least 30x. The WGS data was processed using an in-house Illumina analysis pipeline (https://github.com/UMCUGenetics/IAP). Briefly, reads were mapped to the CRCh37/hg19 human reference genome using BWA-0.7.5a using “BWA-MEM-t 12 -c 100 -M -R” [69]. GATK IndelRealigner [70] was used to realign the reads. Duplicated reads were removed using Sambamba markdup [71].

### Structural variant calling and filtering

Raw SV candidates were called with Manta v0.29.5 using standard settings [72] and Delly v0.7.2 [73] using the following settings: “-q 1 -s 9 -m 13 -u 5”. Only Manta calls overlapping with breakpoint junctions called by Delly (+/− 100 basepairs) were selected. Rare SVs were selected by filtering against SV calls of 1000 Genomes [74] and against an in-house database containing raw Manta SV calls of ~120 samples (https://github.com/UMCUGenetics/vcf-explorer). *De novo* SVs were identified in individuals P1 to P22 by filtering the SVs of the children against the Manta calls (+/− 100 basepairs) of the father and the mother. Filtered SV calls were manually inspected in the Integrative Genome Viewer (IGV). *De novo* breakpoint junctions of individuals P1 to P21 were validated by PCR using AmpliTaq gold (Thermo Scientific) under standard cycling conditions and by Sanger sequencing. Primers were designed using Primer3 software (Additional file S2: Table S2). Breakpoint junction coordinates for individuals P22 to P39 were previously validated by PCR [51,53].

### Single nucleotide variant filtering

Single nucleotide variants and indels were called using GATK HaplotypeCaller. For individuals P1 to P21 (whose parents were also sequenced), reads overlapping exons were selected and the Bench NGS Lab platform (Agilent-Cartagenia) was used to detect possible pathogenic *de novo* or recessive variants in the exome. *De novo* variants were only analysed if they affect the protein structure of genes that are intolerant to missense and loss-of-function variants. Only putative protein changing homozygous and compound heterozygous variants with an allele frequency of <0.5% in ExAC [31] were reported.

### RNA-sequencing and analysis

RNA-seq libraries were prepared using TruSeq Stranded Total RNA Library Prep Kit (Illumina) according to the manufacturer’s protocol. RNA-seq libraries were pooled and sequenced on a NextSeq500 (Illumina) in 2×75bp paired-end mode. Processing of RNA sequencing data was performed using a custom in-house pipeline (https://github.com/UMCUGenetics/RNASeq). Briefly, reads were aligned to the CRCh37/hg19 human reference genome using STAR 2.4.2a [75]. The number of reads mapping to genes were counted using HTSeq-count 0.6.1 [76]. Genes overlapping with SV breakpoints (eg truncated genes) were also analyzed separately by counting the number of reads mapping to exons per truncated gene fragment (up- and downstream of the breakpoint junction). RNA-seq data obtained from PBMCs (Individuals P1 to P22) and LCL cell lines (Individuals P23 to P39) were processed as separate datasets. The R-package DESeq2 was used to normalize raw read counts and to perform differential gene expression analysis for both datasets separately [77]. Genes with more than 0.5 reads per kilobase per million mapped reads (RPKM) were considered to be expressed.

### Gene annotation

Gene information (including genomic positions, Ensembl IDs, HGNC symbols and Refseq IDs) was obtained from Ensembl (GRCh37) using the R-package biomaRt (v2.38) [78]. Genes containing a RefSeq mRNA ID and a HGNC symbol were considered as protein-coding genes. Genomic coordinates for the longest transcript were used if genes contained multiple RefSeq mRNA IDs. The list of 19,300 protein-coding genes was further annotated with 1) pLI, 2) RVIS, 3) haploinsufficiency (HI) scores, 4) OMIM identifiers and 5) DD2GP information for each gene (see Additional file 2: Table S5 for data sources). A phenotypic score based on these five categories was determined for each gene. Modes of inheritance for each gene were retrieved from the HPO and DD2GP databases.

### Computational prediction of the effects of SVs on genes

For each patient, the protein-coding genes located at or adjacent (< 2Mb) to the *de novo* SV breakpoint junctions were selected from the list of genes generated as described above. The HPO terms linked to these genes in the HPO database [19–21] were matched to each individual HPO term assigned to the patient and to the combination of the patient’s HPO terms using Phenomatch [16,17]. For each gene, the number of phenomatchScores higher than 5 (“phenomatches”) with each individual HPO term of a patient was calculated. The strength of the association (none, weak, medium or strong) of each selected gene with the phenotype of the patient was determined based on the total phenomatchScore, the number of phenomatches, the mode of inheritance and the phenotypic score (Fig. S4a, Table 1).

Subsequently, potential direct and indirect effects of the SVs (none, weak or strong) on the genes were predicted (Fig. S4a, Table 1). The prediction analyses were based on chromatin organization and epigenetic datasets of many different cell types obtained from previous studies (see Additional file 2: Table S5 for data sources).

First, we determined which TADS of 20 different cell types overlapped with the *de novo* SVs and which genes were located within these disrupted TADs [37–39] (Additional file 1: Fig. S4b). To determine if the disrupted portions of the TADs contained regulatory elements that may be relevant for the genes located in the affected TADs, we selected the three cell types in which the gene is highest expressed based on RNA-seq data from the Encode/Roadmap projects [45] reanalysed by Schmitt et al [37] (Additional file 1: Fig S4C). The number of active enhancers (determined by chromHMM analysis of Encode/Roadmap ChIP-seq data [45]) in the TADs up- and downstream of the breakpoint junction in the three selected cell types was counted (Additional file 1: Fig S4D). Virtual 4C was performed by selecting the rows of the normalized Hi-C matrices containing the transcription start site coordinates of the genes. The v4C profiles were overlapped with the breakpoint junctions to determine the portion of interrupted Hi-C interactions of the gene (Additional file 1: Fig. S4e). In addition, promoter capture Hi-C data of 22 tissue types [40–43] and DNAse-hypersensitivity site (DHS) connections [44] were overlapped with the SV breakpoints to predict disruption of long range interactions over the breakpoint junctions (Additional file 1: Fig. S4f).Genes with at least a weak phenotype association and a weak SV effect are considered as T3 candidate genes. Genes were classified as T1 candidate drivers if they have a strong association with the phenotype and are strongly affected by the SV. Genes classified as T2 candidate driver can have a weak/medium phenotype association combined with a strong SV effect or they can have a medium/strong phenotype association with a weak SV effect (Fig. 2a, Table 1).

### SV and phenotype information large patient cohorts

Breakpoint junction information and HPO terms for 228 individuals (excluding the individuals already included in this study for WGS and RNA-seq analysis) with mostly balanced SVs were obtained from Redin et al [51]. Phenotype and genomic information for 154 patients with *de novo* copy number variants ascertained by clinical genomic arrays were obtained from an inhouse patient database from the University Medical Center Utrecht (the Netherlands).

## List of abbreviations

HPO: Human Phenotype Ontology
kb: kilobase
RPKM: reads per kilobase per million mapped reads
SNV: Single nucleotide variant
SV: Structural variant
TAD: topologically associated domain
VUS: Variant of unknown significance
WGS: Whole-genome sequencing

## Declarations

### Ethics approval and consent to participate

All individuals or their parents provided written informed consent to participate in the study. The study was approved by the Medical Ethics Committee (METC) of the University Medical Center Utrecht (NL55260.041.15 15-736/M). The study was performed in accordance with the Declaration of Helsinki.

### Consent for publication

All participants in this study provided consent for publication.

### Availability of data and material

Whole genome sequencing and RNA sequencing datasets generated during the study have been deposited in the European Genome-phenome Archive (https://www.ebi.ac.uk/ega)accession number EGAS00001003489. All custom code used in this study is available on https://github.com/UMCUGenetics/Complex_SVs.

## Authors’ contributions

SM and JV performed experiments and computational analyses. JG and RH ascertained and enrolled individuals P1 to P21 and provided phenotypic information. JK and SM cultured LCL cell lines. NB and LdlF performed DNA and RNA isolations and lab support. SB, RJ, MJvR and WK provided support for computational analyses. EvB collected genomic and phenotypic information of individuals U1-U111. GP provided material for individual P22. MET provided LCL cell lines and information of individuals P22-P39. SM, JV, JG, JK, and EC designed the study. SM, JV and EC wrote the manuscript. All authors read and approved the final manuscript.

## Acknowledgements

We wish to thank all individuals who participated in this research. We thank Giulia Pregno and Giorgia Mandrile for the clinical and biological study of individual P22. We thank Utrecht Sequencing Facility (USEQ) for providing RNA-sequencing service and data. USEQ is subsidized by the University Medical Center Utrecht, Hubrecht Institute and Utrecht University. We would also like to thank the Hartwig Medical Foundation for providing whole genome sequencing services.

## Additional files

**Additional file 1 (pdf):** Figures S1 to S8, including figure legends and supplemental references.

**Additional file 2 (xlsx): Table S1.** Phenotype information of the 39 included patients with *de novo* SVs. **Table S2.** Coordinates of the *de novo* SV breakpoint junctions detected in the 39 individuals by WGS. **Table S3.** Candidate driver genes detected for each included patient. **Table S4.** Fusion genes detected in the patients by RNA sequencing. **Table S5.** List of external data sources used in this study. **Table S6.** Detected *de novo* copy number variants in 154 patients of the diagnostics cohort. **Table S7**. Candidate driver genes detected in patients’ cohorts.

## Supplemental materials

**Fig. S1.**
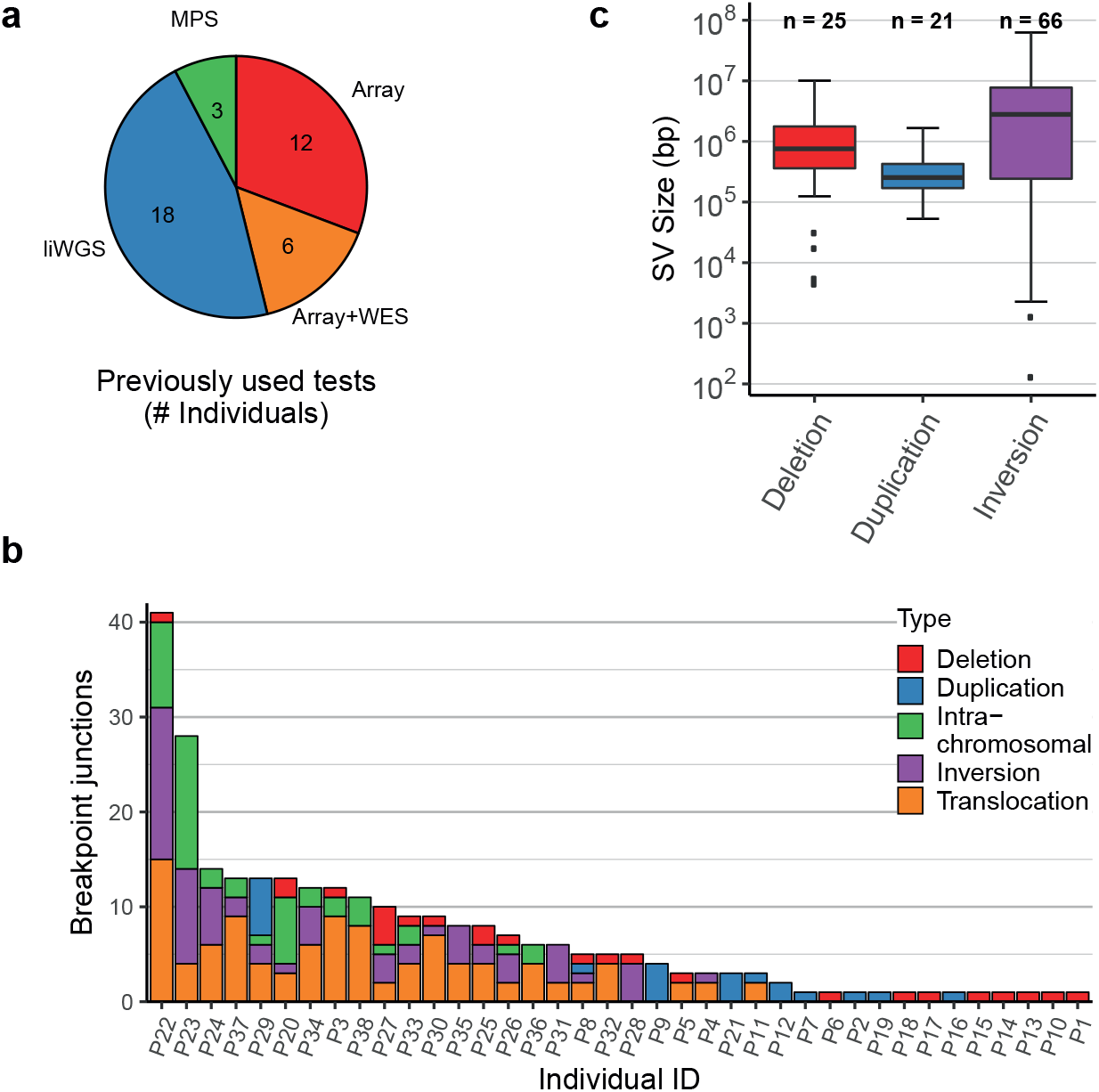
Detected *de novo* germline SVs in 39 included patients. **a** Genetic tests previously used in a clinical setting to identify the *de novo* SVs in the included individuals. Microarrays (ArrayCGH or SNP arrays) were used to detect the deletions and duplications in 18 of the included individuals. MPS: Mate-pair sequencing, WES: Whole Exome Sequencing, liWGS: long-insert Whole Genome Sequencing. **b** Number of identified *de novo* SV breakpoint junctions per individual. **c** Size distribution in base pairs (bp) of the identified *de novo* deletions (median size 757,378 bp), duplications (median size 253,729 bp) and inversions (median size 2,295,988 bp).

**Fig. S2.**
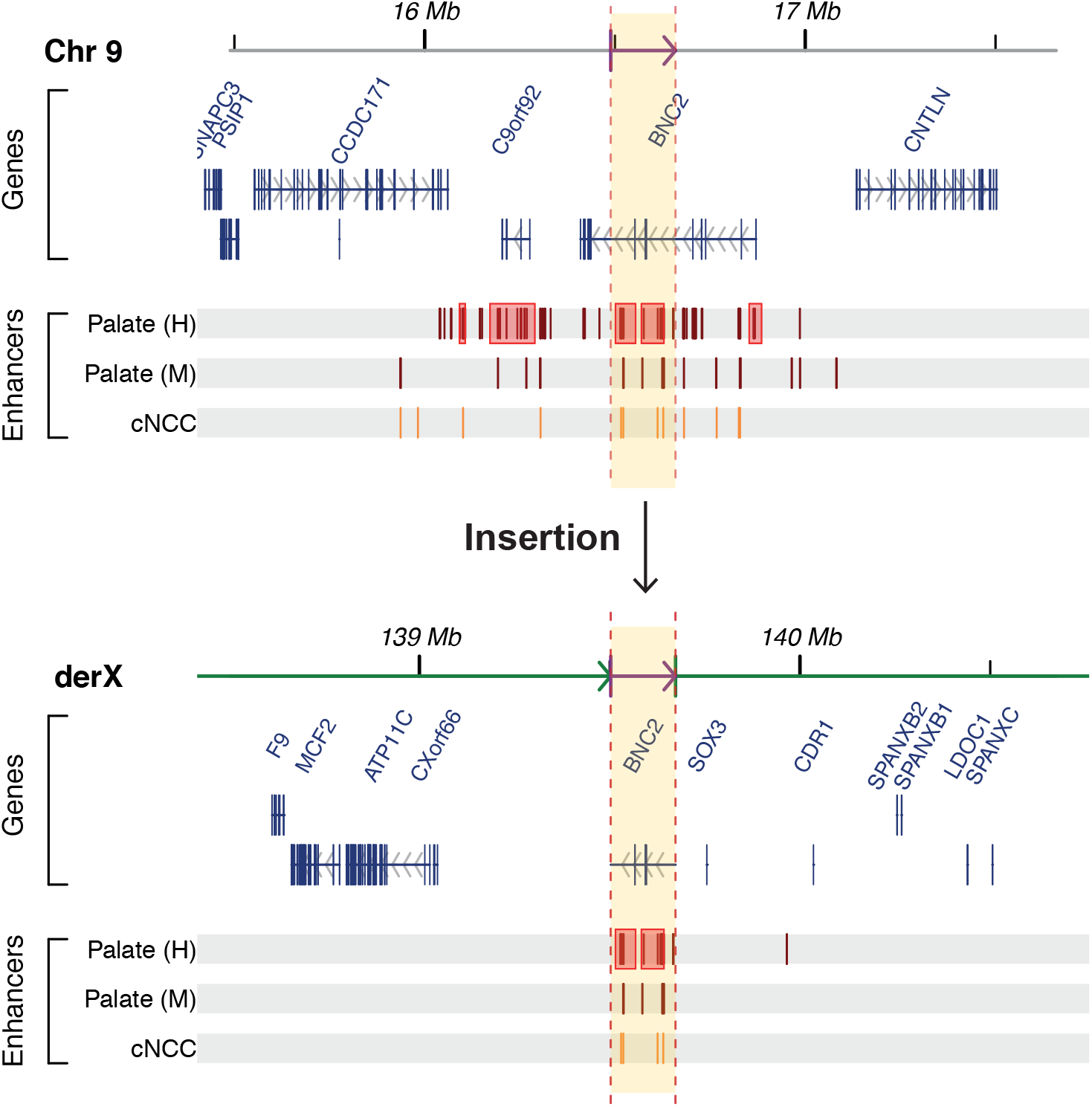
Insertion of a super-enhancer region upstream of *SOX3* detected by WGS in individual P11. A 170kb duplication in the *BNC2* gene body at chr9 was reported by array-based analysis (top panel), but WGS detected that this duplication is actually inserted in chrX (bottom panel). The fragment (highlighted in yellow) is inserted 82 kb upstream of the *SOX3* gene. This locus at chrX contains a palindromic sequence that is susceptible for formation of genomic rearrangement. Multiple patients with varying phenotypes and different insertions at this locus have been described [1–5]. The inserted fragment from chr9 contains multiple enhancers, including two previously described super-enhancer clusters (highlighted by red boxes), that are active in human (Palate(H), Carnegie stage 13) and mouse (Palate (M), embryonic day 11.5) craniofacial development and human cultured cranial neural crest cells (cNCC) [6–8]. The inserted enhancers may disturb the normal expression of the *SOX3* gene and/ or the surrounding genes, which may have led to the cleft palate phenotype in this patient. Genomic coordinates of mouse (mm9) embryonic craniofacial enhancers (determined by p300 ChIP-seq [6]) were converted to hg19 coordinates using LiftOver (https://genome.ucsc.edu/cgi-bin/hgLiftOver).

**Fig. S3.**
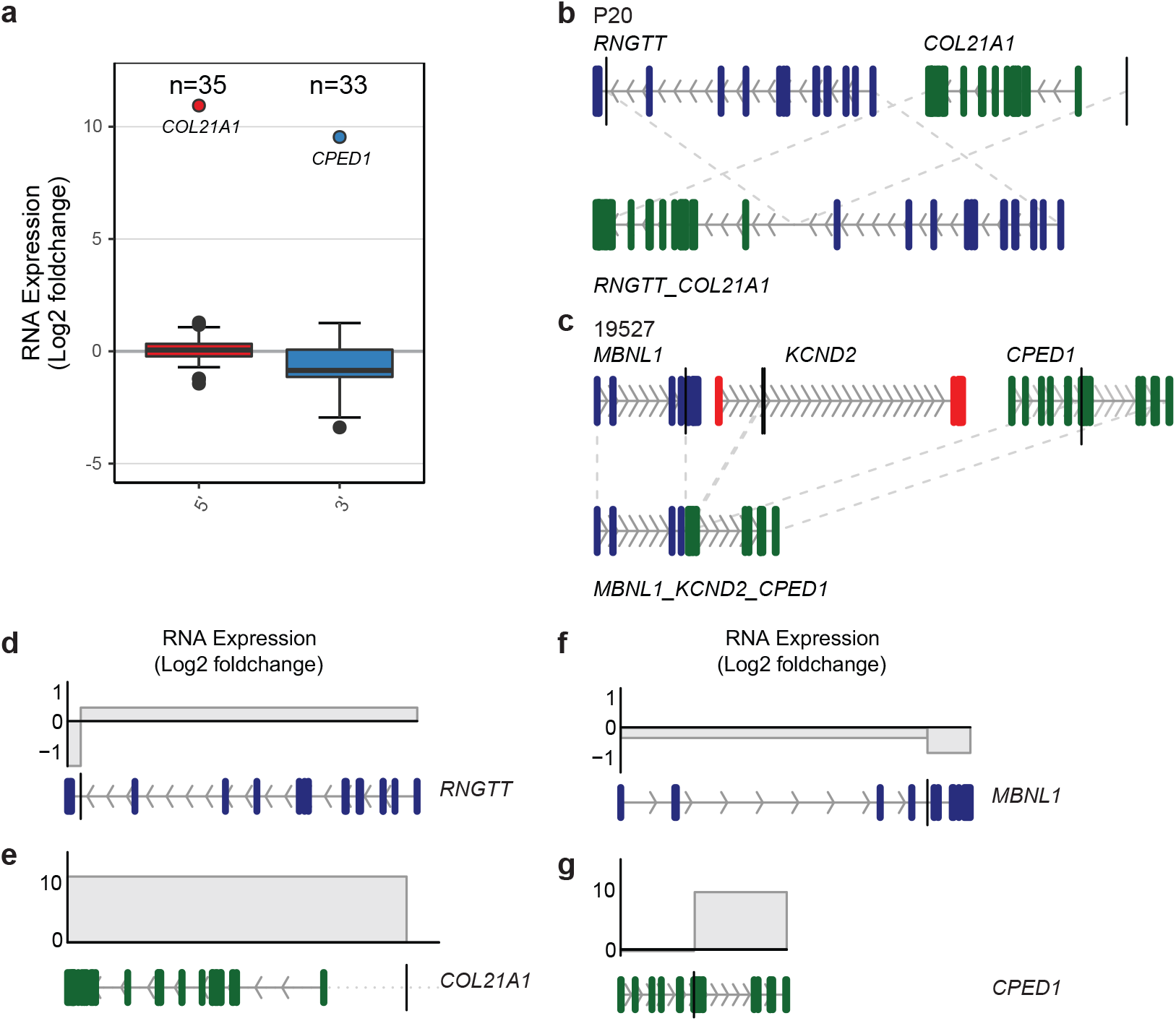
RNA expression of genes truncated by *de novo* germline SVs. **a** Log2 fold change expression values (compared to expression of the exons in control individuals) for 5’ gene fragments and 3’ gene fragments of truncated genes. The 5’ fragment of *COL21A1* and the 3’ fragment of *CPED1* show a strong overexpression due to a gene fusion. **b** Schematic representation of the *RNGT-T_COL21A1* fusion gene caused by genomic rearrangements in individual P20. The breakpoint junctions near the *RNGTT* (ENST00000369485) and *COL21A1* (ENST00000244728) gene bodies are depicted by the vertical black lines. **c** Schematic reconstruction of the *MBNL1_KCND2_CPED1* fusion gene in individual P34. Breakpoint junctions in the truncated genes *MBNL1* (ENST00000324210),*KCND2*(ENST00000331113)and *CPED1*(ENST00000310396) are represented by the vertical black lines. **d - g** RNA log2 fold change expression values (compared to the expression in unaffected individuals) for the fragments of the truncated genes *RNGTT*, *COL21A1*, *MLBNL1* and *CPED1*.

**Fig. S4.**
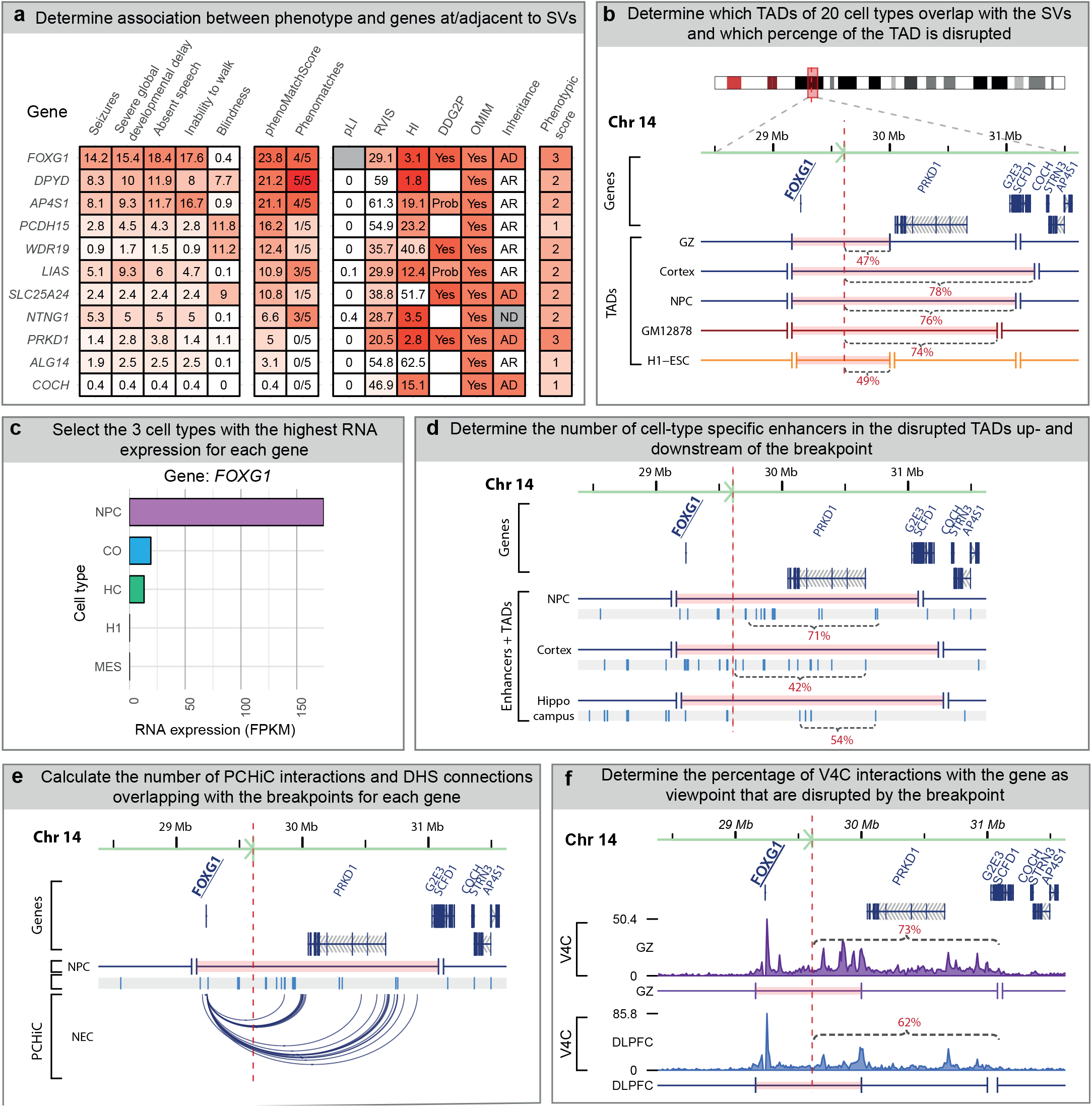
Schematic overview of computational strategy used to predict positional effects. **a** The association of a gene at or adjacent to an SV with the patient-specific phenotype is based on the Phenomatch score, the number of phenomatches, mode of inheritance and the phenotypic score. Each gene has a fixed phenotypic score (ranging from 1 to 5) based on it spLI (> 0.9), RVIS (<10) and haploinsufficiency (HI,<10) scores and the presence of the gene in DDG2P and OMIM. **b** The TADs of 20 different cell types are overlapped with the SV breakpoint junctions of the individual. The TADs affected by a breakpoint are split into fragments up- and down-stream of the breakpoint and the size of each fragment (relative to the size of the intact TAD) is calculated. Subsequently the genes located on each fragment are determined and each gene receives a score based on the relative size of the fragment. For example, 47% of the TAD in germinal zone (GZ) cells containing *FOXG1* is considered disrupted. **c** For each gene the 3 cell types with the highest RNA expression (FPKM: Fragments Per Kilobase Million) based on the Encode/Roadmap ChIP-seq data are selected. **d** For each gene, enhancers from the three selected cell types are overlapped with the disrupted TAD fragments. The number of enhancers in the disrupted part of the TAD is compared to the number of enhancers in the TAD fragment containing the gene. This ratio is considered at the percentage of enhancers moved away from the gene (for example, the location of 71% of neural progenitor cell enhancers in the *FOXG1* TAD is changed). **e** For each gene, PCHiC interactions of 22 cell types and promoter-DHS connections are overlapped with the breakpoint junctions. The number of interactions overlapping with the junctions is divided by the total number of interactions of the gene. For example, for *FOXG1* all 107PCHiC interactions (in multiple cell types) overlap with the breakpoint junction. **f** Virtual 4C profiles were generated for each gene and these were overlapped with the breakpoint junctions to determine the percentage of interactions that are located up- or downstream of the breakpoint junction. For *FOXG1*, 73% of the V4C interactions in dorsolateral prefrontal cortex (DLPFC) cells are considered to be disrupted.

**Fig. S5.**
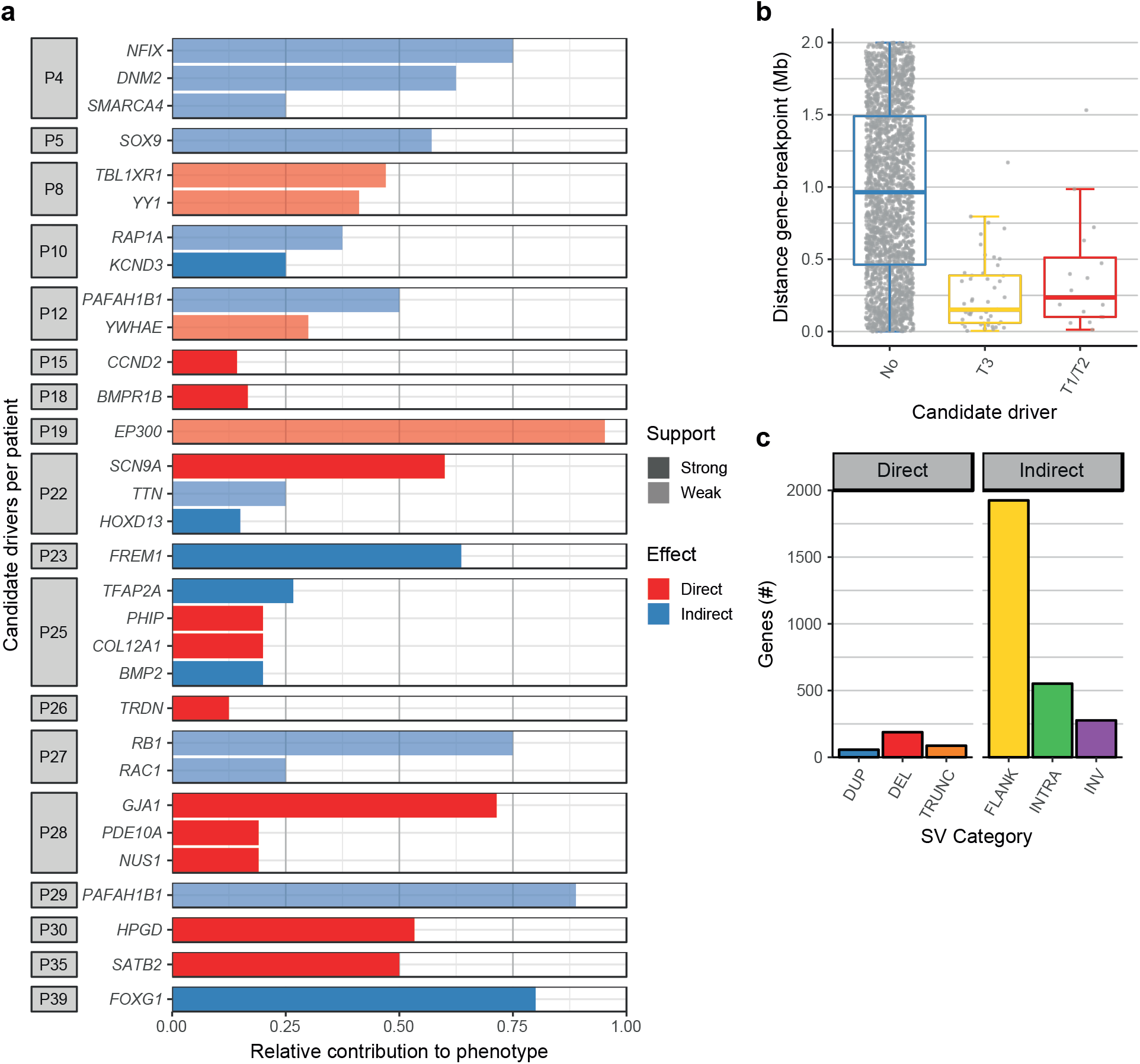
Overview of the detected candidate driver genes. **a** Relative contributions of the candidate drivers the phenotypes of the individuals. The contributions are based on the number of Phenomatch hits (pheomatch Score > 5) of a gene with each individual HPO term assigned to an individual, e.g. a gene with a contribution of 0.75 is associated with 75% of the HPO terms of an individual. Shading indicates if there is relatively weak or strong evidence for an effect on the candidate driver. **b** Genomic distance (in base pairs) between the indirectly affected candidate driver genes (adjacent to the SVs) and the closest breakpoint junction. Most predicted candidate drivers are located within 1 Mb of a breakpoint junction. **c** Total number of analysed genes per SV category. DUP: Duplication, DEL: Deletion, TRUNC: Truncation, FLANK: Flanking region (+/− 2Mb), Intra: Intrachromosomal rearrangement, INV: Inversion.

**Fig. S6.**
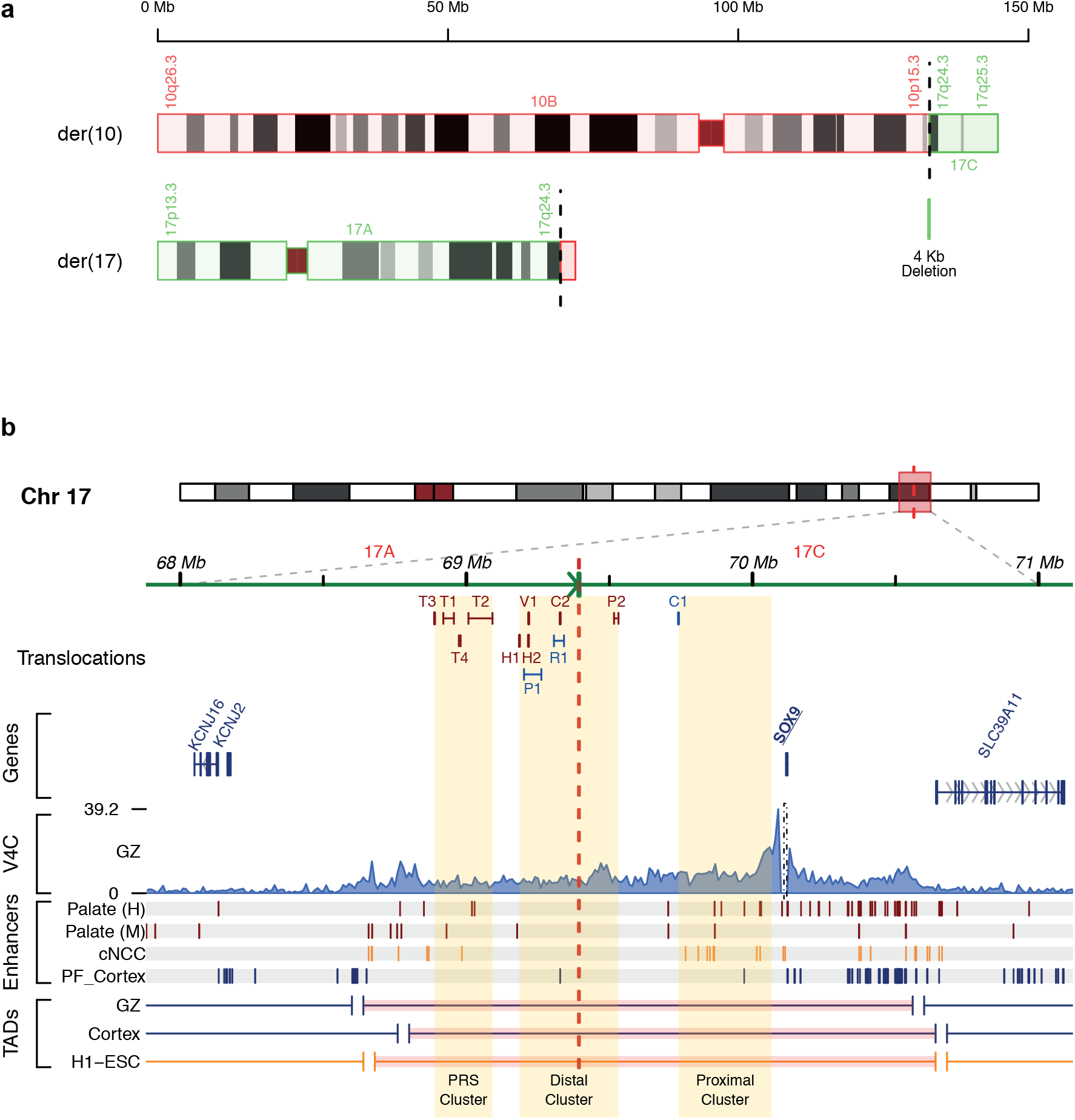
Prediction of positional effects of a translocation on SOX9 in individual P5. **a** Ideogram of the derivative chromosomes in individual P5. WGS identified a *de novo* translocation between chromosome 10 and 17 (46,XY,t(10;17)(p15;q24)). The breakpoint on chr17 (chr17:69395684, indicated by the vertical dotted red line) is 721 kb upstream of *SOX9*. A small 4kb fragment from chr17 is deleted (chr17:69391279-69395683). **b** Genome browser overview showing region surrounding the translocation breakpoint (red dotted line) at chromosome 17 in individual P5. The phenotype of this individual is characterized by acampomelic campomelic dysplasia and Pierre-Robin Syndrome including cleft palate, micrognathia and a long philtrum. SVs including translocations have been detected upstream of *SOX9* in individuals with various phenotypes including campomelic dysplasia. The translocations found in patients with phenotypes including cleft palate are shown in red and translocations found in patients with different phenotypes are depicted in blue. These translocations are predicted to separate *SOX9* from enhancers active in the developing palate, which may lead to the cleft palate phenotypes. Information about the other patients was obtained from the following publications: T1+T2+T3 [9]; T4 [10]; C1+C2 [11]; P1+P2 [12]; V1 [13]; R1 [14]; H1+H2[15].

**Fig. S7.**
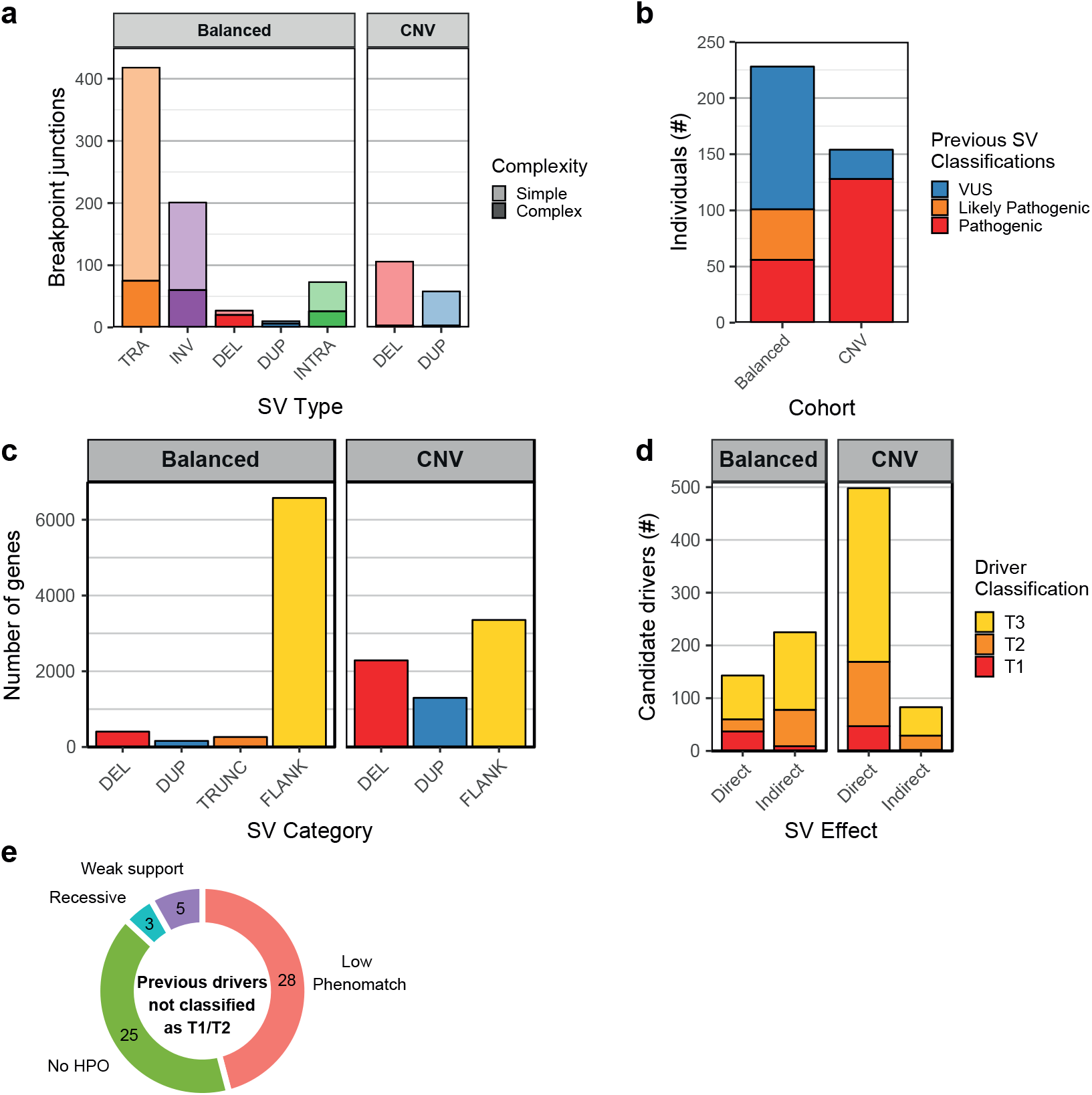
Overview of SVs and candidate drivers in two cohorts of patients with *denovo* SVs. **a** Quantification of previously identified *de novo* SVs in a cohort containing patients with mostly balanced SVs and a cohort containing patients with copy number variants (CNV). *De novo* translocations (TRA), inversions (INV) and intra-chromosomal rearrangements (INTRA) are most prevalent in the cohort of patients with balanced SVs. Some patients have complex genomic rearrangements (>3SVs)including some deletions (DEL) or duplications (DUP). The cohort labelled as “CNV” consist of patients with relatively simple deletions and duplications (<10 Mb in size). **b** Number of patients whose *denovo* SVs were previously classified as pathogenic, likely pathogenic or variant of unknown significance (VUS) per cohort. **c** Total number of analysed genes per SV category in the two cohorts. Dup: Duplicated, Del:Deleted, Trunc:Truncated, Flank: Flanking SVs (<1Mb). **d** Total number of predicted directly and indirectly affected candidate drivers per cohort. **e** Quantification of the genes that were previously classified as pathogenic or likely pathogenic(by Redin et al[16]), but not identified as T1 or T2 candidate driver by our approach. These classification differences may be caused by a lack of HPO terms associated with the gene, low phenomatch scores below the threshold of our method, insufficient (weak) support for an effect of an SV on the gene detected by our method or a presumed recessive mode of inheritance.

**Fig. S8.**
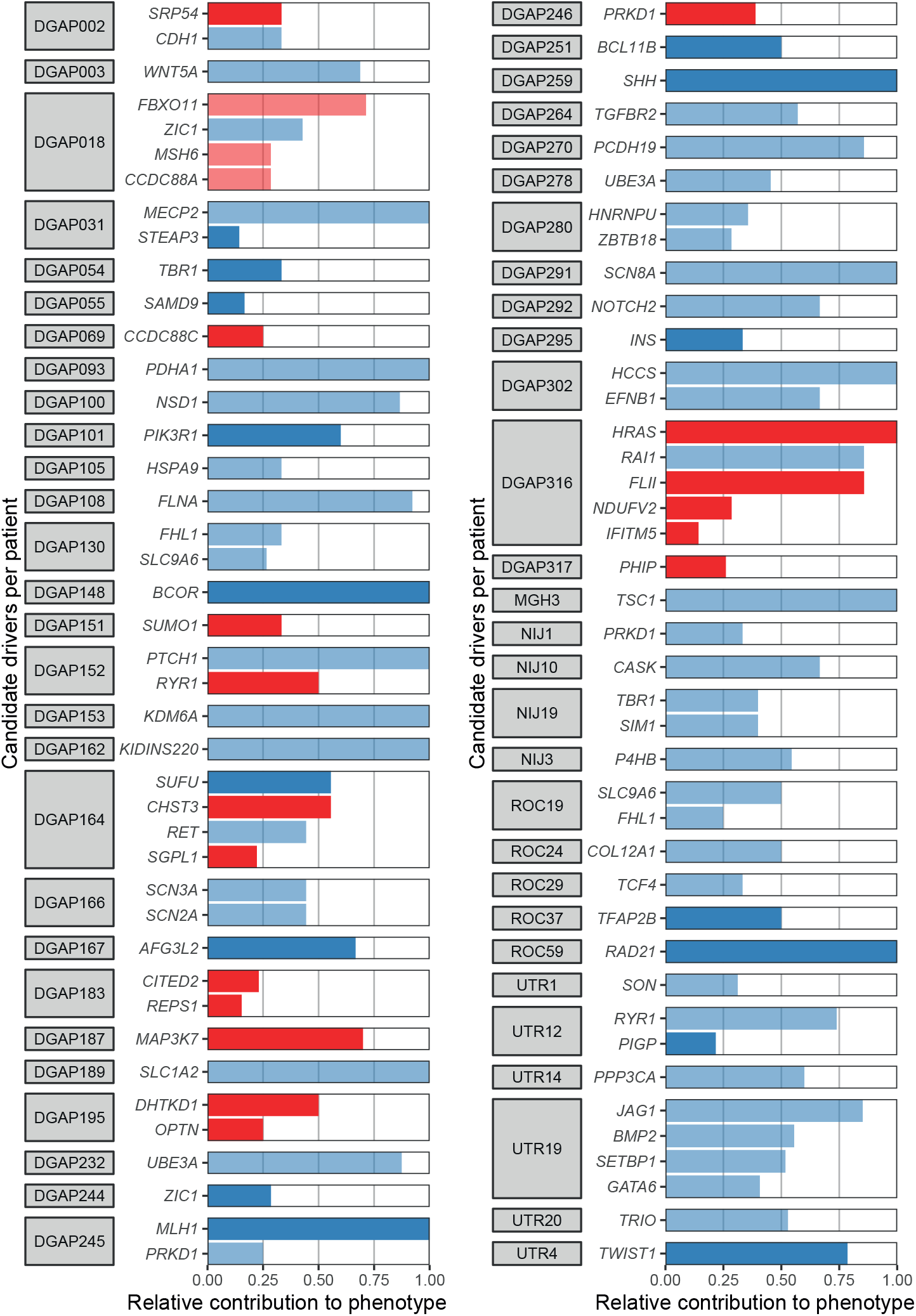
Predicted contributions of candidate drivers to the phenotypes of patients with balanced structural variants of unknown significance. T1/T2 candidate drivers were detected in 31 patients whose *de novo* SVs were previously classified as VUS by Redin et al [16]. The contributions to the phenotypes are based on the number of Phenomatch hits (phenomatchScore>5) of the gene with each individual HPO term used to describe the phenotype of a patient. Shading indicates if there is relatively weak or strong evidence for an effect on the candidate driver.

